# Identification of green lineage osmotic stress pathways

**DOI:** 10.1101/2021.07.19.453009

**Authors:** Josep Vilarrasa-Blasi, Tamara Vellosillo, Robert E. Jinkerson, Friedrich Fauser, Tingting Xiang, Benjamin B. Minkoff, Lianyong Wang, Kiril Kniazev, Michael Guzman, Jacqueline Osaki, Michael R. Sussman, Martin C. Jonikas, José R. Dinneny

## Abstract

Maintenance of water homeostasis is a fundamental cellular process required by all living organisms. Here, we use the green alga *Chlamydomonas reinhardtii* to establish a foundational understanding of evolutionarily conserved osmotic-stress signaling pathways in the green lineage through transcriptomics, phosphoproteomics, and functional genomics approaches. Five genes acting across diverse cellular pathways were found to be essential for osmotic-stress tolerance in Chlamydomonas including cytoskeletal organization, potassium transport, vesicle trafficking, mitogen-activated protein kinase and chloroplast signaling. We show that homologs of these genes in the multicellular land plant *Arabidopsis thaliana* have conserved functional roles in stress tolerance and reveal a novel PROFILIN-dependent actin remodeling stage of acclimation that ensures cell survival and tissue integrity upon osmotic stress. This study highlights the conservation of the stress response in algae and land plants and establishes Chlamydomonas as a unicellular plant model system to dissect the osmotic stress signaling pathway.

---

Maintaining cellular water homeostasis is vital in all living organisms, requiring various mechanisms to counteract differences in water availability between the intracellular and extracellular environment occurring during fluctuating environmental conditions^1^. Detailed mechanistic insight of the osmotic stress response has been established in unicellular model systems such as yeast, but knowledge is more limited in photosynthetic organisms^2^. We sought to advance our understanding by establishing the unicellular alga Chlamydomonas (Chlamydomonas reinhardtii) as a model system to study osmotic stress through a systems biology approach and identify functionally conserved components across the green lineage.

## Chlamydomonas transcriptomic and post-translational response to osmotic stress

To establish a temporal map of the osmotic stress responses in Chlamydomonas, we performed a transcriptomic analysis of the wild-type strain CC-4533 across a 24-hour time course using two osmotic challenges, NaCl and mannitol, treatments that substantially reduce cell growth but are non-lethal (Extended Data Fig. 1a-b). We identified a total of 1,456 differentially-regulated genes (FC >2, FDR <0.01), with a peak of regulation after 1 hour (Table S1, Table S2, Extended Data Fig. 1c-d). Up-regulated genes were enriched (FDR < 0.05) in annotations associated with cellular stimulus, vesicle-mediated-transport, and amyloplast organization while downregulated genes were associated with the cell cycle, rRNA metabolic processes, chloroplast fission, and deoxyribonucleotide metabolism (Fig. 1a). Consistent with these data, we found that an osmotic challenge induced an increase in the number of vesicles and starch puncta in Chlamydomonas cells (Fig. 1b, Extended Data Fig. 2a-b), using fluorescent reporters that mark these compartments^3^. These regulatory patterns suggest that osmotic stress induces a pause in general cellular functions and activates stress response genes, a pattern also observed in yeast and land plants upon osmotic stress^4,5^. The Chlamydomonas genome includes homologous gene families found in both Saccharomyces (*Saccharomyces cerevisiae*, fungi), and Arabidopsis (*Arabidopsis thaliana*, land plant)^6^. Chlamydomonas osmotic stress-responsive genes have homologs in both Arabidopsis and Saccharomyces, although the likelihood of differential expression upon osmotic stress is greater in Arabidopsis homologs compared to Saccharomyces homologs (24% and 6% respectively) (Material and methods, Extended data Fig. 3, Table S2). Our analysis therefore suggests that, although the Chlamydomonas transcriptional response to osmotic stress shares features with both Arabidopsis and Saccharomyces, it is more similar to Arabidopsis.

**Figure 1.**
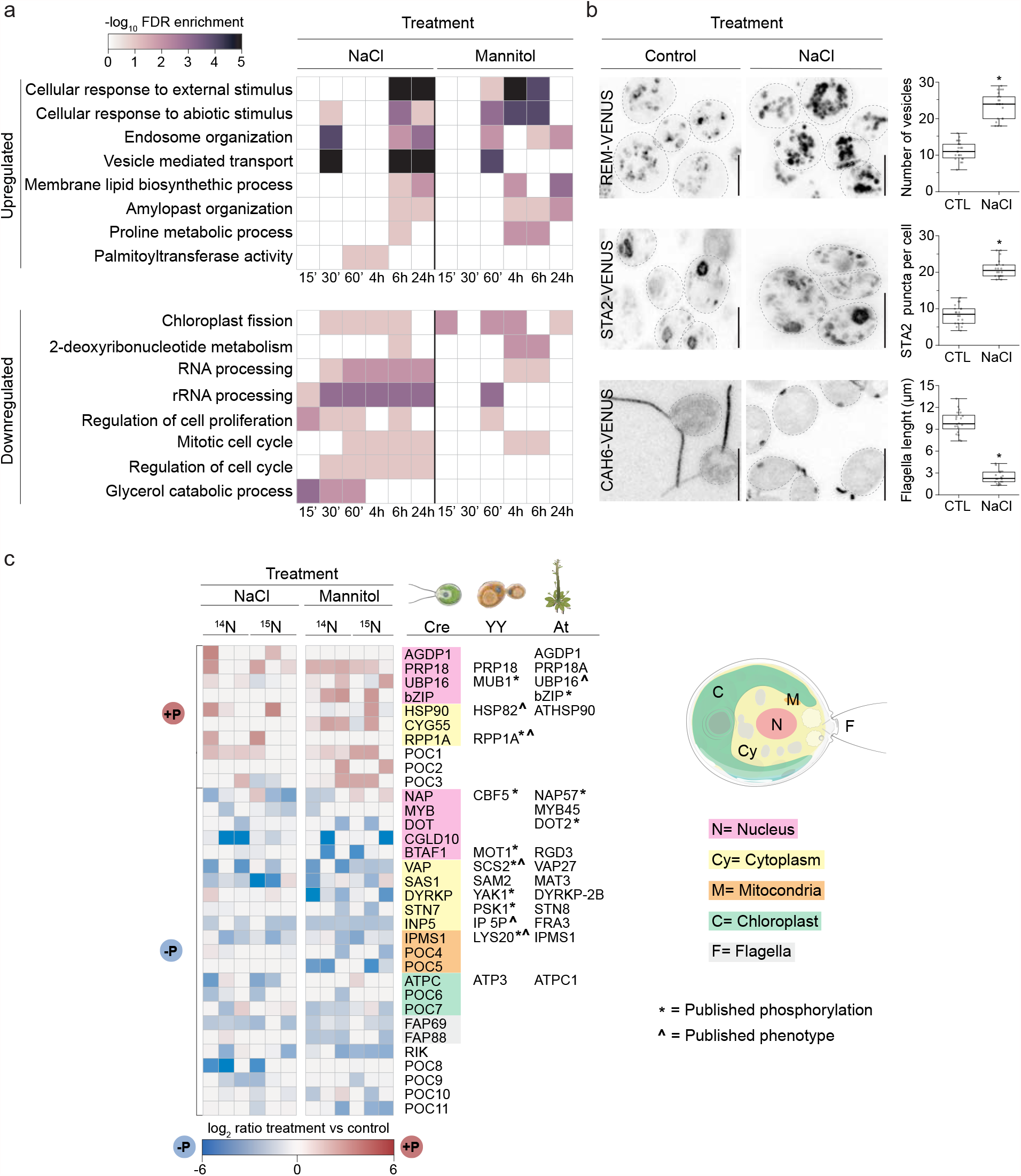
*Chlamydomonas reinhardtii* shared and divergent osmotic stress responses with other eukaryotes. **a**, Gene ontology term analysis of the transcriptional response of Chlamydomonas cells treated with 100 mM NaCl or 300 mM mannitol at the indicated time points, ‘ and h indicate minutes and hours respectively (See Extended Data 1-3, Table S1, S2). **b**, Confocal images of fluorescently tagged Chlamydomonas cells with different fluorescent markers^3^ under control conditions and treatment with 100 mM NaCl for 30 hours. Cellular markers used: Endosomes, Cre16.g661700: REMORIN, labels the Golgi apparatus and secretory pathway (REM-VENUS); Starch, Cre17.g721500: STARCH SYNTHASE 2 (STA2), labels pyrenoid starch and starch granules (STA2-VENUS); flagella, Cre12.g485050: CARBONIC ANHYDRASE 6 (CAH6) labels flagella (CAH6-VENUS). (*Right*) Quantification of the number of vesicles in REMORIN cells, STA2 puncta and flagella length in control and cells treated with NaCl. n > 20 cells per condition. Scale bars = 10 μm. **c**, Heat map representing proteins differentially phosphorylated (FC >2, FDR <0.01) upon 5 min treatment with 100 mM NaCl or 300 mM mannitol. Color code represents protein localization according to PredAlgo^40^ or associated functions. * Represents published phosphorylated proteins upon osmotic stress, **^** represents proteins with a described role in osmotic pathways (See Table S2, S3 and Extended Data 3).

While the abscisic acid (ABA) hormone signaling pathway controls much of the transcriptional response to osmotic stress in Arabidopsis, in Chlamydomonas, we did not find enrichment of gene ontologies associated with either ABA regulated genes or related cis-elements (eg. ABA RESPONSE ELEMENT or G-box) (Extended Data Fig. 1e-f)^7,8^. This, in conjunction with the lack of orthologous genes to ABA receptors, or other downstream components, in the Chlamydomonas genome^6^, suggests the absence of a functional ABA-mediated transcriptional response to osmotic stress in this species. A feature of Chlamydomonas cells that is shared with animals and certain plant lineages is the formation of motile cilia. We found an enrichment of genes encoding ciliary proteins being differentially expressed in Chlamydomonas under osmotic stress^9^ (Fisher’s Exact Test=1E-05) (Extended Data Fig. 2c, Table S2), as well as a substantial reduction in flagellar length upon osmotic stress (Fig. 1b). Together, these data demonstrate that osmotic stress induces rapid changes in Chlamydomonas cellular organization that are likely under the control of signaling pathways distinct from the dominant ABA-dependent pathway of land plants.

To uncover early signaling components of the Chlamydomonas osmotic stress response pathway, we adapted untargeted high-resolution mass spectrometry in conjunction with metabolic labeling to examine the effect of 5 min NaCl or mannitol treatment (Fig. 1c, Table S3-S4). We identified 33 Chlamydomonas differentially phosphorylated proteins (FC >2, FDR <0.01) encompassing a variety of functions including: transcriptional regulation (MYB and bZIP family proteins) and flagellar machinery (Flagella-associated proteins (FAP), FAP 69 and FAP 88), among others. We found several proteins with homologs that when mutated in Arabidopsis have osmotic sensitive phenotypes, such as; Ubiquitin protease UBIQUITIN-SPECIFIC PROTEASE 16 (UBP16)^10^ and ENHANCED EM LEVEL (EEL), a bZIP transcription factor homologous to ABA-INSENSITIVE 5 (ABI5)^11,12^. Similarly, we identified several proteins with Saccharomyces homologs previously shown to be involved in osmotic stress, including; ribosomal stalk protein RIBONUCLEASE P (RPP1A)^13^, VESICLE-ASSOCIATED MEMBRANE PROTEIN-ASSOCIATED PROTEIN (SCS2)^14^, and INOSITOL POLYPHOSPHATE 5-PHOSPHATASE (IP 5-P), a Phosphoinositol binding protein involved in cytoskeletal reorganization^15^. Among the Saccharomyces orthologous proteins, 8 out of 14 were shown to be phosphorylated upon osmotic stress, while only 3 out of 16 showed phosphorylation in Arabidopsis (Fig. 1c, Table S4). We identified 56 proteins with orthologs that are also phosphorylated in Arabidopsis upon osmotic stress with lower confidence (FC >1.5, FDR <0.01) (Extended Data Fig. 4, Table S4). Our dataset includes 11 unannotated proteins which we renamed PHOSPHORYLATED UPON OSMOTIC STRESS IN CHLAMYDOMONAS (POC).

Altogether our transcriptional and phosphoproteomic analysis established the immediate targets of osmotic stress signaling in Chlamydomonas and highlights components most likely to be conserved across the green lineage and with non-photosynthetic eukaryotes.

### Discovery of novel genes with roles in osmotic stress

Microbial model systems provide advantages for genome-wide, forward genetic screens due to the ease of culturing thousands of genotypes simultaneously. We utilized a recently developed barcoded genome-wide mutant library^16^ and subjected Chlamydomonas cells to various osmotic stresses. We used four different perturbations: NaCl, mannitol, Polyethylene glycol (PEG), and hypo-osmotic stresses and identified 76 genes with a growth defect when mutated across at least one stress: 27 for NaCl, 34 for mannitol, 13 for hypoosmotic, 2 for PEG (FDR<0.3; Fig. 2a-c, Table S5-6). While several replicates were performed for each condition, few significant mutants were identified across replicates, suggesting that the screens had not been performed to saturation. We validated our mutant phenotypes by performing secondary screens of 140 mutants selected according to different statistical criteria (Table S7, Extended Data Fig. 5), validating 55% of the hits. We identified genes encoding: 9 transporters, 4 flagella-associated proteins, 7 kinases, 2 phosphatases, 11 nuclear localized proteins, among others. From this list of high confidence hits we identified 34 and 53 Saccharomyces and Arabidopsis orthologs, respectively (FDR< 0.3, Table S5). Interestingly, 24 genes did not have any previous annotation in Chlamydomonas, Arabidopsis, or Saccharomyces, and therefore we renamed them <u>OSMO</u>TIC GROWTH DEFECTIVE IN CHLAMYDOMONAS (OSMO). Among our top hits, we identified HYPEROSMOLALITY-INDUCED Ca INCREASE 1 (CreOSCA1) and MscS-like (CreMSL) that mediate the initial events of osmotic stress signaling in Arabidopsis^17,18^. Both showed sensitivity to NaCl and mannitol, acclimation to osmotic stress and flagella defects, and cellular localization comparable to the Arabidopsis homolog (Extended Data Fig. 6)^19,20^. Below, we describe several examples dissecting our functional genomics results by cellular functions.

**Figure 2.**
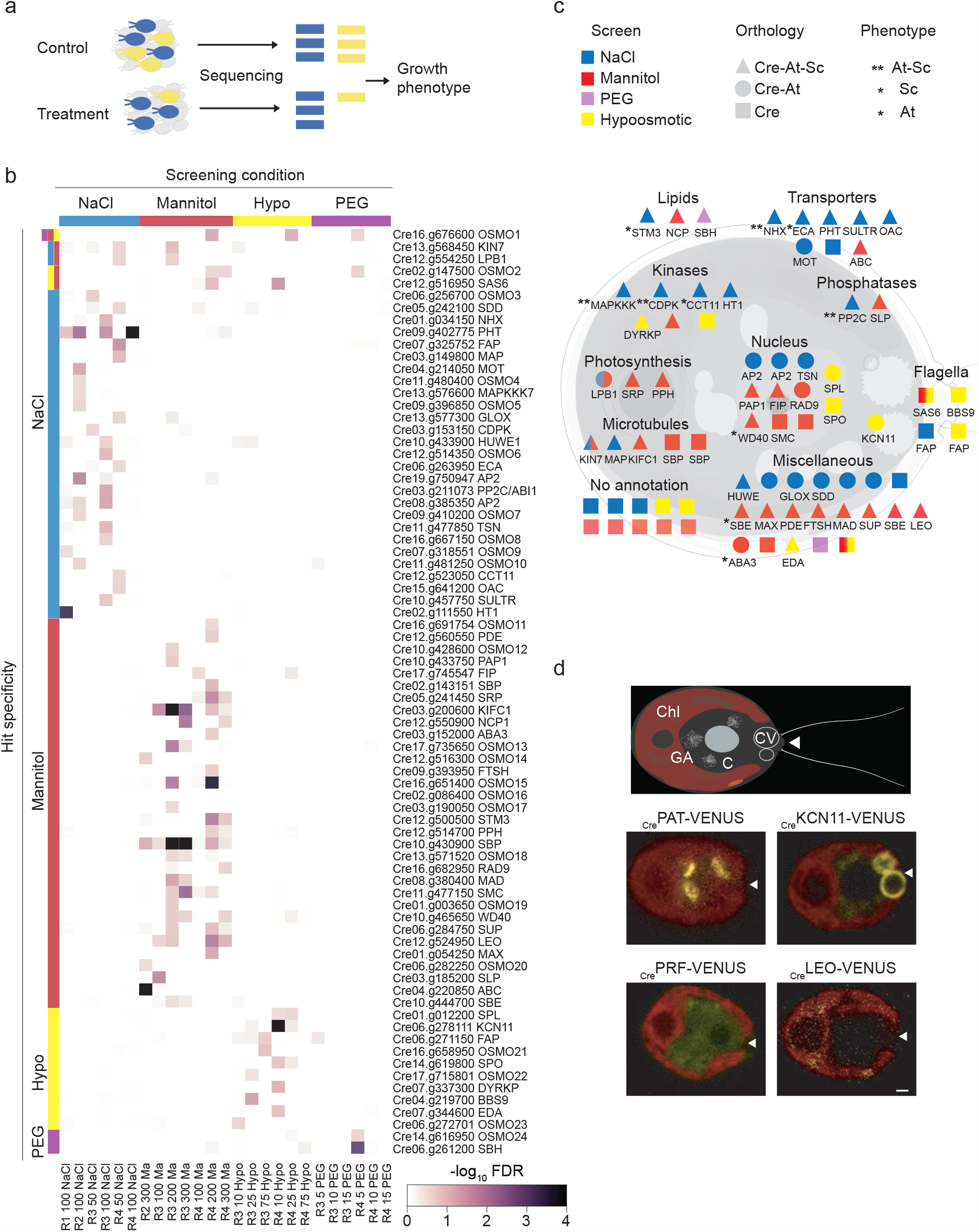
Chlamydomonas osmotic stress pathways involve different organelles. **a**, Diagrammatic representation of barcode mutant screens. Unique barcodes allow genome-wide screening of Chlamydomonas mutants in a pool. Mutants sensitive to osmotic stress can be identified because their barcodes will be less abundant after growth under treatment (NaCl, mannitol, PEG, hypoosmotic) compared to control conditions. **b**, Heat map representing significance of hits (>2 alleles per gene, see Material and Methods). **c**, Diagrammatic representation of high confidence hits from genome-wide barcoded screens upon different osmotic stresses (FDR <0.3). Hit orthologs to *S. cerevisiae* (Sc) or *A. thaliana* (At) based on reciprocal BLAST (< 1E-10). Osmotic phenotype represents hits with a previously described role in osmotic stress in *S. cerevisiae* (Sc.), *A. thaliana* (At) or both (See Table S5). **d**, Confocal images of Chlamydomonas cells showing the subcellular localization of proteins fused to VENUS fluorescent protein. Cartoon represents different organelles in Chlamydomonas: Chl = chloroplast, GA = Golgi apparatus, C = cytoplasm, CV = contractile vacuole. Red color shows chloroplast autofluorescence and yellow shows VENUS fluorescence. Scale bar = 1 μm.

We found that core osmotic stress signaling components such as kinases: CALCIUM-DEPENDENT PROTEIN KINASE (CPDK) and MITOGEN-ACTIVATED PROTEIN KINASE KINASE KINASE (MAPKKK) homologous to Cre03.g153150 and Cre13.g576600, and phosphatases: PROTEIN PHOSPHATASE 2C (PP2C) and SHEWENELLA-LIKE PROTEIN PHOSPHATASE (SLP), homologous to Cre03.g211073 and Cre03.g185200, have conserved functions between kingdoms, playing central roles in Arabidopsis and Saccharomyces osmoregulatory pathways^1,2^(Fig. 2c, Table S6). Our screens have osmolyte specificity as shown by the number of membrane transporters necessary for Chlamydomonas growth under NaCl, but not other osmotic stresses, including SODIUM/HYDROGEN EXCHANGER (NHX, Cre01.g034150) and ENDOPLASMIC RETICULUM-TYPE CALCIUM-TRANSPORTING ATPASE (ECA, Cre06.g263950) transporters, which have been previously shown to play a role in sodium detoxification in plants and fungi^21,22^ and other transporters such as a SULFATE TRANSPORTER (SULTR, Cre10.g457750) and a PHOSPHATE TRANSPORTER (PHT, Cre09.g402775).

Nuclear localized factors identified in sensitive mutants included: transcription factors with homology to the APETALLA-2 family (AP2, Cre19.g750947 and Cre08.g385350) and SQUAMOSA PROMOTER PROTEIN-LIKE (SPL, Cre03.g185200); a WD-40 repeat-containing protein, Cre10.g465650, previously linked to drought stress in Arabidopsis^23^; and a core component of the homologous recombination pathway RADIATION DEFECTIVE (RAD, Cre16.g682950), among others. Interestingly we found two genes whose Arabidopsis homologs were previously shown to function in acclimation to osmotic stress: a STARCH BRANCHING ENZYME (SBE, Cre10.g444700) and ABA DEFICIENT 3 (ABA3, Cre03.g152000) a gene encoding a molybdenum cofactor sulfurase that has been demonstrated to participate in both ABA and ABA-independent stress pathways within Arabidopsis^24,25^. Not surprisingly, we also found limited overlap in the gene lists identified through transcriptomics and functional genomics, with just five genes exhibiting a significant growth defect and transcriptional response upon osmotic stress; an AP2 transcription factor, SBE, a motor microtubule protein KINESIN 7 (KIN7, Cre13.g568450), a proprotein convertase subtilisin (Cre05.g242100) and Cre17.g735650, which is not annotated. Overall, our functional genomics identified new high confidence genes involved in osmotic stress signaling and corroborates low overlap between transcriptomics and functional genomics, previously reported for other model systems, such as yeast^26,27^.

Our systems biology approach points to the involvement of multiple organelles in the osmotic response pathway. We picked genes with two validated mutant alleles from our secondary screens and studied the localization of their encoded proteins fused to VENUS (Fig. 2d, Extended Data Fig. 7). We were able to localize four proteins to various cellular organelles, including a Potassium channel (KCN11, Cre06.g278111) at the contractile vacuole, S-palmitotransferase (PAT, Cre06.g277000) at the Golgi apparatus, <u>L</u>ETHAL <u>E</u>MBRYONIC <u>O</u>SMOTIC (LEO, Cre12.g524950) at the chloroplast, and <u>PRO</u>FILIN (PRO, Cre10.g427250) in the cytoplasm and at the nuclear envelope. Together our multi-omics characterization of Chlamydomonas points to a conservation of many osmotic signaling pathway components between algae, fungi, and land plants and identifies new components with high confidence for future studies.

### Novel osmotic stress pathways conserved across the green lineage

Previous studies have found that using a combination of model systems is advantageous for functionally characterizing conserved gene functions, such as using yeast and mouse to dissect the cell cycle, or Chlamydomonas and Arabidopsis to uncover the photosynthetic apparatus^28^. Thus, to identify new osmotic cellular pathways conserved across the green lineage, we grew Arabidopsis seedlings with mutations in genes homologous to Chlamydomonas osmo-sensitive genes identified in our screens and analyzed the ability of their roots to acclimate to osmotic stress induced by NaCl or mannitol containing media. We were able to uncover five genes with mutants exhibiting osmotic phenotypes in both Chlamydomonas and Arabidopsis; including genes encoding for: MITOGEN ACTIVATED PROTEIN KINASE KINASE (MAPKK), GATED OUTWARDLY-RECTIFYING POTASSIUM CHANNEL (GORK), S-PALMITOTRANSFERASE (PAT), PROFILIN 5 (PRF-5), and LETHAL EMBRYONIC OSMOTIC 1 (LEO1), a gene that encodes an uncharacterized putative GTP-binding protein (Fig. 3a, Extended data Fig. 5d).

**Figure 3.**
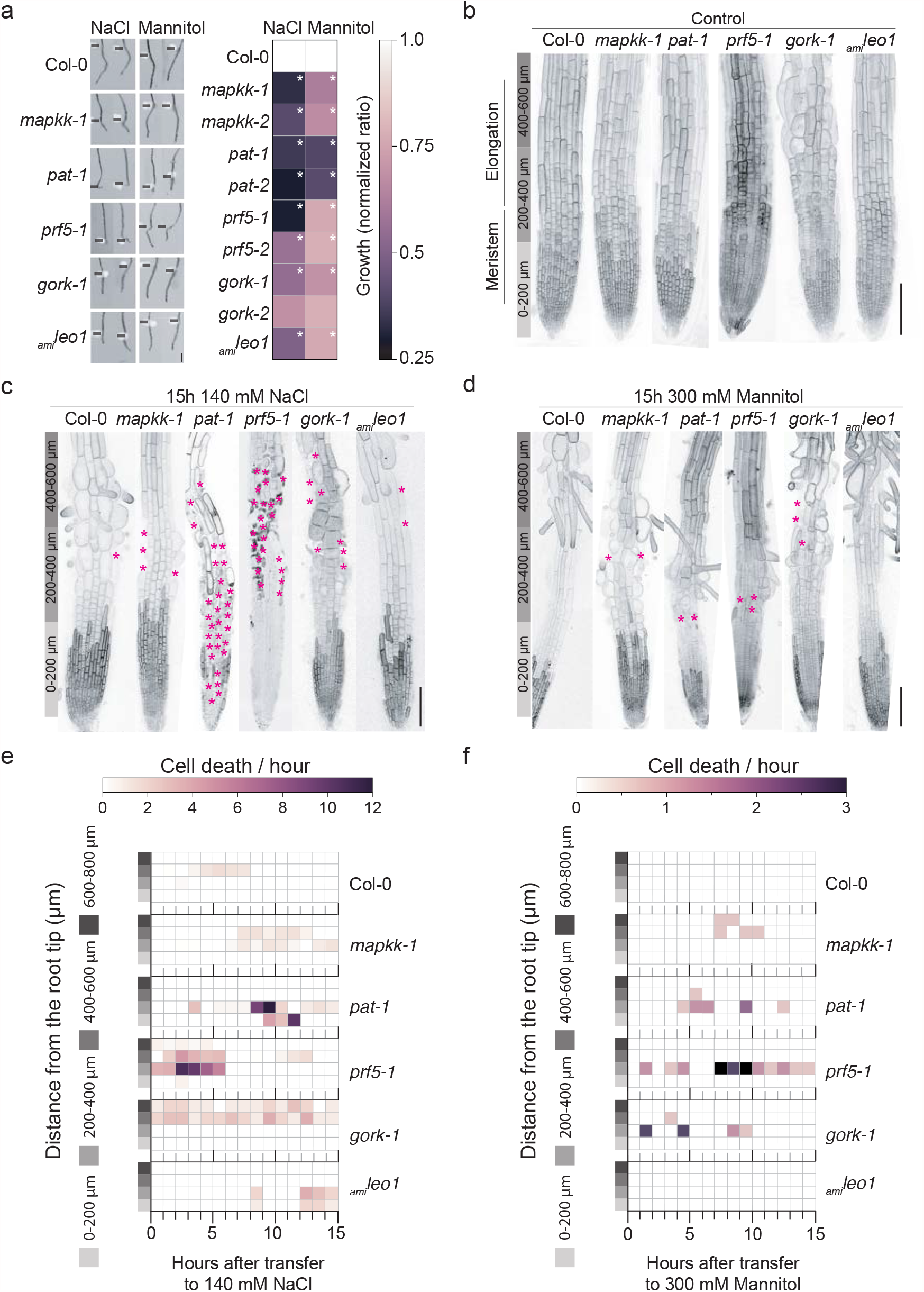
Conserved osmotic stress pathways show different spatiotemporal cell death patterns in Arabidopsis roots. **a**, (*Left*) *A. thaliana* primary roots 4 days post-transfer to 140 mM NaCl or 300 mM mannitol, black lines indicate transfer side. Scale bar = 1 mm. (*Right*) Quantification of root growth after transfer to 140 mM NaCl or 300 mM mannitol, root growth ratio upon transfer was normalized to the growth of wild type upon stress treatment. * Indicates *p*-value < 0.05, two-way ANOVA. **b**, Confocal images of 4 days post germination primary roots, plasma membrane marked by LTI6b-YFP fluorescence as in C and D. Scale bar = 100 μm. **c**, Confocal images of 4 days post germination root tips, 15 hours after transfer to 140 mM NaCl. * Indicates cell death. Scale bar = 100 μm. **d**, Confocal images of 4 days post germination root tips, 15 hours after transfer to 300 mM mannitol. * Indicates cell death. Scale bar = 100 μm. **e**, Quantification of root cell death after transfer to 140 mM NaCl. n > 10 roots for each genotype. **f**, Quantification of root cell death after transfer to 300 mM mannitol. n > 10 roots for each genotype.

Time-lapse imaging of the Arabidopsis primary roots with an introgressed plasma membrane marker, *LTI6b-YFP*^*29*^, revealed different patterns of root growth cessation upon osmotic stress and the cellular basis for these growth defects in the mutants (Fig. 3b-f). Under standard conditions, all mutant primary roots grew similar to wild-type except *gork-1*, which exhibited a loss of root hair anisotropic growth, producing round hair cells. Inflated root hairs were suppressed when the mutants were grown on either media lacking supplemented potassium, at low temperatures, or by removing sucrose from the media, suggesting that the abnormal growth and loss of cellular integrity in this mutant is caused by a growth-dependent intracellular accumulation of potassium (Extended data Fig. 8).

Wild-type plants transferred to 140 mM NaCl showed cell death in the late elongation and early differentiation zones, approximately 400-600 µm from the root tip within 4-8 hours after transferring to stress. In comparison, treatment with 300 mM mannitol did not affect cell viability. In contrast, we could score different spatiotemporal cell-viability phenotypes in our mutants (Fig. 3c-f). The *mapkk-1* mutant showed cell death in the elongation-differentiation zone upon NaCl or mannitol treatment, but with a peak of damage to the root occurring at different times post-treatment (7-14 hours for NaCl, 7-10 hours for mannitol). *Pat-1* mutants showed cell death in the meristematic zone, 0-200 µm from the root tip, and elongation zone with a peak of cell death occurring 9 hours after transfer to stress treatment, which resulted in roots without epidermal cells within the first 400 µm from the root tip. Similarly, mannitol treatment induced cell death to a minor degree in *pat-1* mutants, starting 4 hours after transfer. Early cell death occurred in the meristematic and elongation zones of *prf-5* mutants upon transfer to NaCl, leaving most epidermal cells dead. Comparably, mannitol treatment promoted cell death at the meristematic-elongation zone boundary (a.k.a. transition zone) as early as 2 hours after treatment in *prf-5* mutants. Continuous cell death in the transition zone occurred in *gork-1* mutants after NaCl treatment, while mannitol promoted cell death in the early elongation zone. Finally, cell death occurred in the *leo-1* mutant at late timepoints upon NaCl treatment, while no cell death happened upon mannitol treatment. Together our phenotypic characterization indicates that all mutants exhibit sensitivity to both NaCl and mannitol-mediated osmotic stress, though greater sensitivity was generally observed with NaCl. Mutants exhibited distinct patterns of cell viability defects, which suggests different cellular mechanisms may be mediating the sensitivity to osmotic stress.

### The actin cytoskeleton reorganizes upon osmotic stress

The Arabidopsis *prf-5* mutant exhibited one of the earliest and most severe defects in cell viability amongst our characterized mutants, with most meristematic and elongating cells dying within the first 5 hours after transfer to stress. Profilin is an actin interacting protein that determines the dynamics of the actin cytoskeleton by controlling the rate of polymerization, bundling, and cable formation^30,31^. We therefore hypothesized that osmotic stress may alter actin cytoskeleton dynamics and that the *prf-5* mutant exhibits hypersensitivity to stress due to the inability to properly control this reorganization. To test this hypothesis we monitored actin dynamics in roots using the actin-binding domain of fimbrin fluorescent reporter, ABD2-GFP reporter^32^. Upon transfer to osmotic stress, both 140 mM NaCl and 300 mM mannitol, root cells in the elongation zone underwent massive actin reorganization (Fig. 4a). An increase in the asymmetric distribution of actin filaments, skewness, revealed an increase in actin bundling together with a decrease in the filament number. Concomitantly, actin filament angle switched from being nearly parallel to the cell longitudinal axis of the cell to being perpendicular^33^ (Fig. 4a-b). Interestingly, visualization of actin dynamics in Chlamydomonas using Lifeact-NeonGreen, showed a similar response to osmotic stress with an increase in skewness (Extended data Fig. 10). Together these data suggest that actin reorganization upon osmotic stress is conserved across the green lineage.

**Figure 4.**
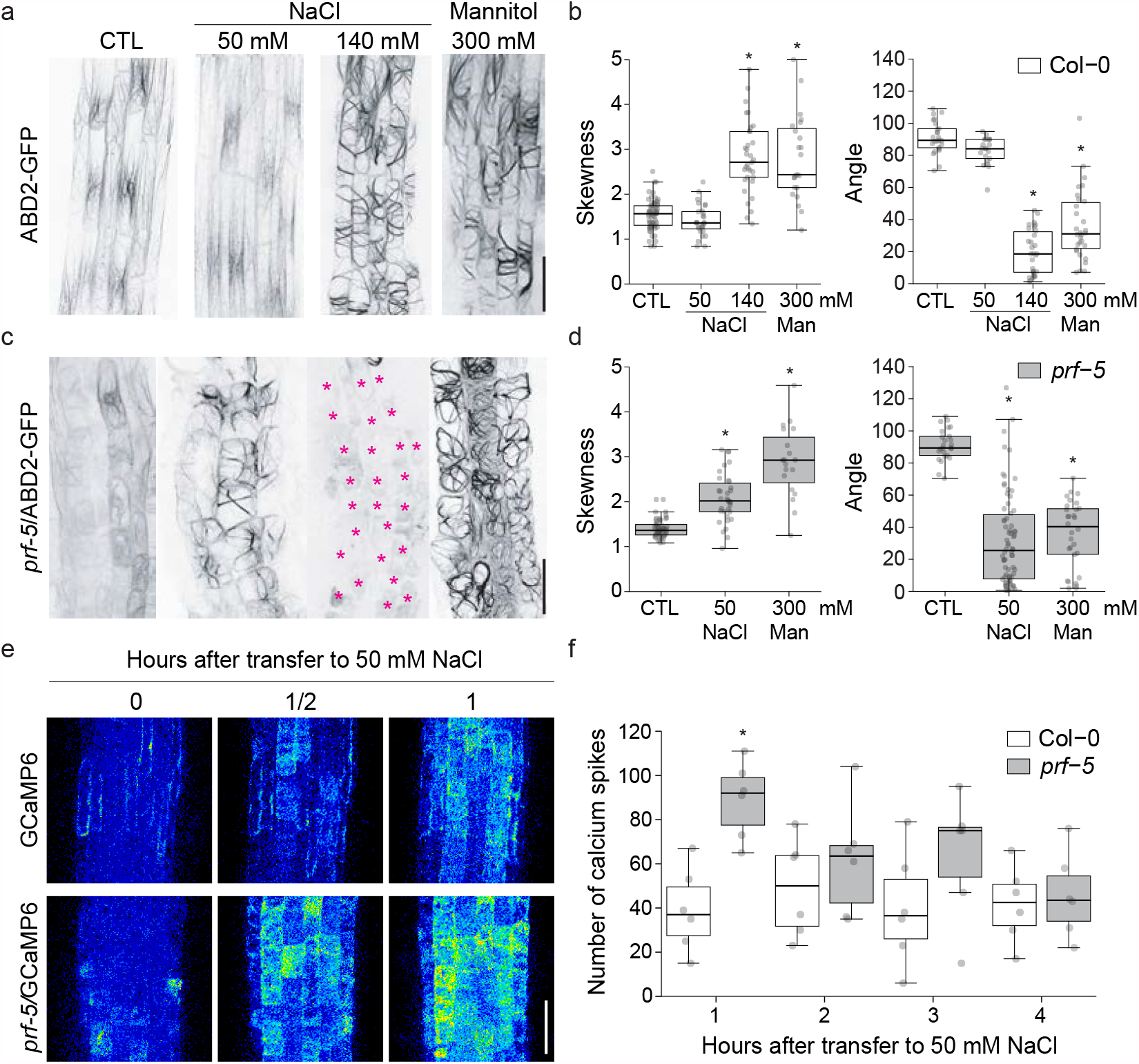
Osmotic stress promotes actin bundling in the root elongation zone. **a**, Actin localization in the Arabidopsis primary root elongation zone of wild-type roots on control conditions (CTL), transferred to 50 mM NaCl, 140 mM NaCl or 300 mM mannitol for 16 hours. Images from treatments correspond to cells actively elongating at the time of transfer. Actin filaments were visualized with the actin binding domain of the FIMBRIN protein tagged with green fluorescent protein (ABD2-GFP). Scale bar = 50 µm. **b**, Quantification of skewness and angle of actin filaments relative to the cell radial axis. Each point represents a cell that was actively elongating when roots were transferred. * Represents *p-value* < 0.05, two-way ANOVA. **c**, Actin localization in the Arabidopsis root elongation zone of *prf-5* roots on control conditions (CTL), transferred to 50 mM NaCl, 140 mM NaCl, or 300 mM mannitol for 16 hours. Images from treatments correspond to cells actively elongating at the time of transfer. Actin filaments visualized with ABD2-GFP. Scale bar = 50 µm. **d**, Quantification of skewness and angle of actin filaments relative to the cell radial axis. Each point represents a cell that was actively elongating when roots were transferred. * Represents p < 0.05, two-way ANOVA. **e**, 16 colors LUT confocal images of primary root elongation zone expressing GCaMP6, and *prf-5/GCamp6*. **f**, Hourly average number of [Ca^2+^] spikes in wild-type and *prf-5* cells expressing GCaMP6 treated with 50 mM NaCl.

Treatment of *prf-5* mutants with 140 mM NaCl led to cell death in the elongation zone (Fig. 3 c, e). Treatments with lower concentrations of NaCl, 50 mM, promoted actin reorganization in the *prf-5* mutant while no changes were observed in wild-type plants, suggesting that the *prf-5* actin network is hypersensitive to osmotic stress (Fig. 4c-d). To localize PRF5 we generated a PRF5 fluorescent tag reporter driven by its own promoter. *ProPRF5::PRF5-YFP* is expressed in root hair cells of the elongation zone and differentiation zone as well as the last two layers of the lateral root cap. Time-course imaging of the expression of PRF5 upon osmotic stress revealed the downregulation of its expression during the initial hours after treatment (Extended data Fig. 10). Since our mutant analysis suggests that PRF5 inhibits NaCl-induced bundling, we hypothesize that PFR5’s normal function may be to delay the induction of osmotic-stress mediated actin bundling. Calcium signaling is an important downstream mediator of abiotic stress^34,35^. To test whether the defects observed in *prf-5* are due to a lack of a stress response or a hyper induction of a stress response, we monitored calcium dynamics in roots transferred to a mild NaCl concentration in wild-type and *prf-5* backgrounds (Fig. 4e-f). Wild-type plants transferred to 50 mM NaCl showed infrequent calcium spikes. In contrast, *prf-5* mutants showed ectopic calcium spikes within the elongation region and an increase in spiking frequency immediately after transfer to stress. Thus, osmotic-stress mediated calcium dynamics are hyperactive in *prf5* mutants, supporting the hypothesis that the dynamical properties of the actin cytoskeleton tune cellular acclimation and sensitivity to changes in water availability.

## Discussion

Our work provides a framework for comparing the conservation and diversification of osmotic stress tolerance pathways. Across Chlamydomonas, yeast and Arabidopsis we have observed common mechanisms to reduce investment in growth and to redirect resources towards osmotic homeostasis. Interestingly, divergence between species occurs frequently in the types of organelles where acclimatory mechanisms function and the downstream signaling pathways that coordinate the response. While Chlamydomonas utilizes flagella as a primary mechanism of exploring the environment, root growth would be the analogous process in seed plants such as Arabidopsis, and both are suppressed under osmotic stress. These data suggest that organisms evolve tolerance mechanisms that suit their specific ecological niche and the physiological processes that they normally carry out.

The actin cytoskeleton is a dynamic network controlling diverse cellular activities such as vesicle trafficking and ion channel activity ^36^. Early cellular responses to osmotic stress involve shifts in the balance between endocytosis and exocytosis of proteins important for growth, such as cellulose synthase complexes ^37–39^, and ion homeostasis, such as ion channels that regulate cell turgor pressure ^18^ and calcium signaling ^19^. We show here that osmotic stress causes a dramatic sequestration of actin into bundles and cables, which may effectively inhibit many of the cellular processes that require filamentous actin. This large scale change in actin organization may provide structural support to the cell or may be part of a quiescence program. Further work is needed to explore the signaling and biophysical implications of this broadly conserved stage of osmotic-stress acclimation.

## Acknowledgments

We thank Xiobo Li for sharing early versions of the mutant library; Silvia Ramundo for generous gift of pRAM118 plasmid; Masayuki Onishi for generously sharing Lifeact-NeonGreen Chlamydomonas strain. Heather Cartwright and the Carnegie imaging facility for microscopy support. Members of the Dinneny lab for helpful discussions. We thank Gregory A Barret-Wilt, and the phosphoproteomic core facility for help and assistance in the generation of the Chlamydomonas osmotic phosphoproteome. We thank Christopher J. Staiger for providing ABD2-GFP lines and Wolf Frommer for providing GcaMP6 reporter.

This project was supported by grants awarded to J.R.D from the NIH NIGMS (R01 GM123259-01) and a Faculty Scholars grant from the Simons Foundation and Howard Hughes Medical Institute (55108515); grants awarded to M.C.J. from NIH (DP2-GM-119137), NSF (MCB-1146621 and MCB-1914989) and the Simons Foundation and HHMI (55108535); grants awarded to M.R.S IOS PGRP No. 2010789 and MCB No. 9143816; Simons Foundation fellowships of the Life Sciences Research Foundation awarded to R.E.J. and J.V.B.; and an EMBO long term fellowship (ALTF 1450-2014) awarded to J.V.B.

## Author contributions

R.E.J., T.X., and J.V.B. analyzed RNA-seq data: BM and MS performed phosphoproteomic experiments; R.E.J., F.F., and J.V.B. prepared mutant pools, performed treatments, and processed all samples, and analyzed data with guidance from in M.C.J.; T.V generated Chlamydomonas fluorescently tagged lines and validated mutant insertion sites; T.V., K.K., and J.V.B. performed and analyzed Chlamydomonas secondary screenings; L.W. generated Cre10.g45568 reporter line; K.K. provided technical assistance and help with the analysis of the phosphoproteomic dataset; M.G. and J.O. performed initial screen of Arabidopsis orthologous phenotypes; J.V.B. performed all the other experiments; J.V.B. and J.D. design the experiments and analyze the data; J.V.B wrote the initial draft of the manuscript; J.D and J.V.B wrote the final version of the manuscript with input from all authors.

## Extended Data Figures and Tables

**Extended Data 1.**
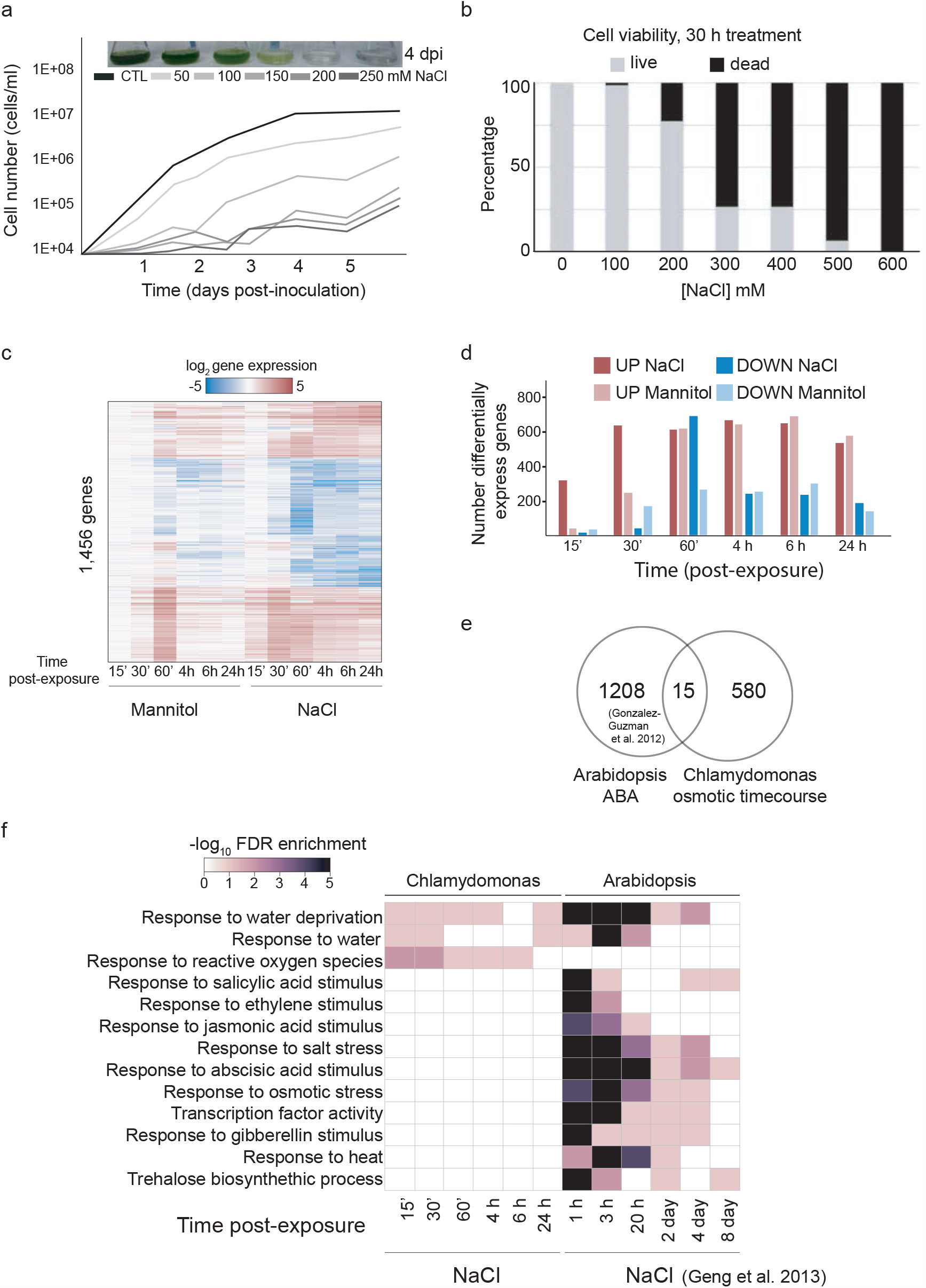
*Chlamydomonas reinhardtii* transcriptional response to osmotic stress lacks obvious signs of ABA hormonal regulation. **a**, Growth curves of Chlamydomonas wild-type cells (CC-4533) grown under different concentrations of NaCl. **b**, Cell viability of Chlamydomonas wild-type cells treated with different concentrations of NaCl for 30 hours. Evans blue staining was used to assess cell viability. **c**, Heat map representing clustered log2 values of 1,456 differentially regulated genes (FC >2, FDR <0.01) under 100 mM NaCl and 300 mM Mannitol treatment at different timepoints. **d**, Number of differentially expressed genes (FC >2, FDR <0.01) at different timepoints. **e**, Overlap of Chlamydomonas differentially expressed genes during NaCl and mannitol time course with *A. thaliana* orthologous and differentially regulated genes upon ABA treatment in *A. thaliana*. (*A. thaliana* ABA regulated genes from Guzman et al 2012). **f**, Gene ontology analysis of differentially regulated genes across the Chlamydomonas time course treated with 100 mM NaCl and NaCl time course of Arabidopsis roots^8^ (See Table S1-S3).

**Extended Data 2.**
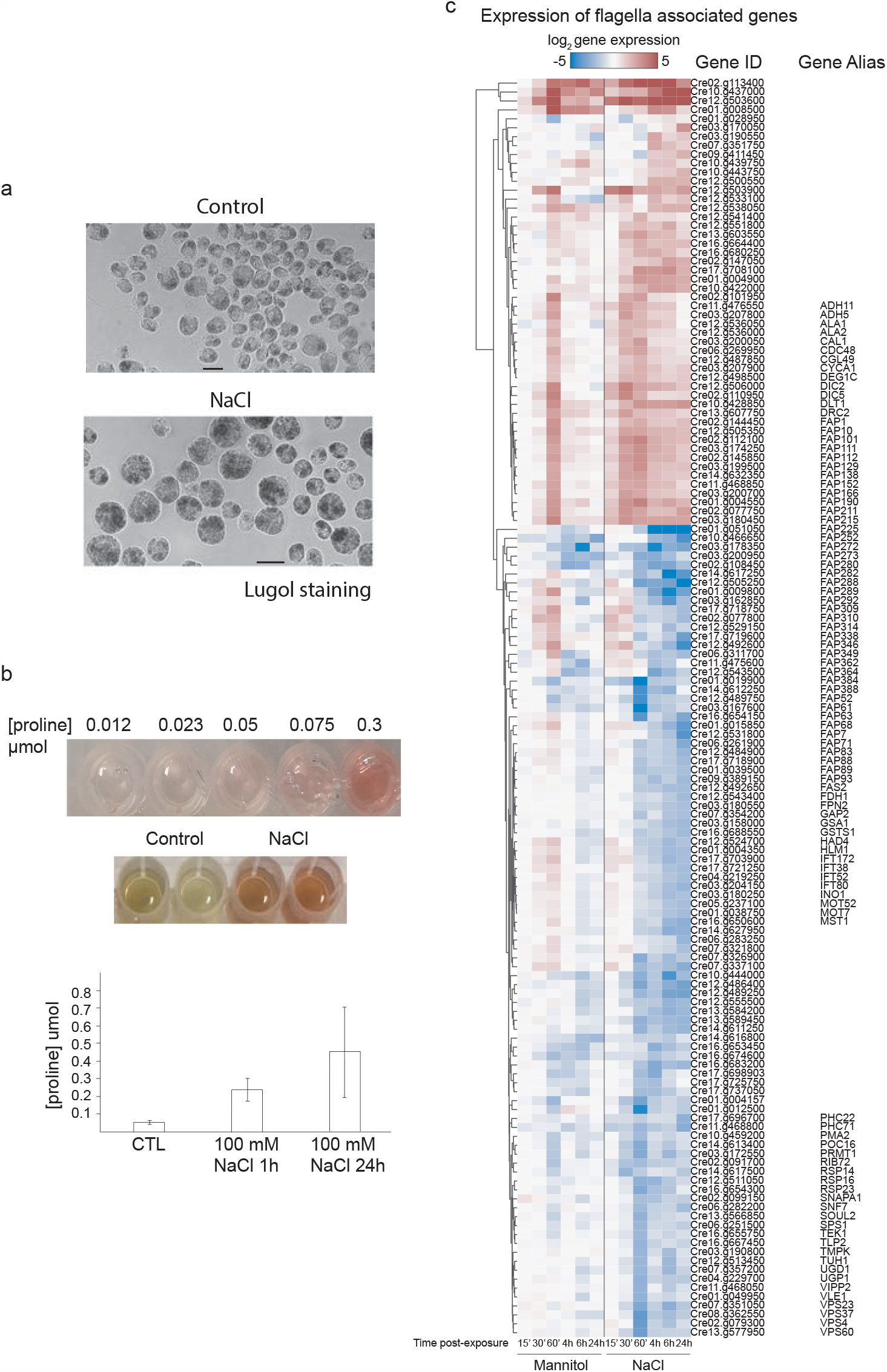
Chlamydomonas osmotic response affects flagella related genes and promotes starch and proline accumulation. **a**, Chlamydomonas cells stained with Lugol. (*Top*) CTL; Control Chlamydomonas cells in mid-early exponential phase stained with Lugol. (*Bottom*) NaCl, Chlamydomonas cells in mid-early exponential phase treated with 100 mM NaCl for 20 hours and stained with Lugol. Scale bars = 10 μm. **b**, Proline quantification of Chlamydomonas cells upon NaCl treatment. **c**, Heat map representing log2 gene expression of Chlamydomonas Ciliary proteins (The Chlamydomonas Flagellar proteome project (http://chlamyfp.org/index.php)) differentially regulated in the Chlamydomonas osmotic transcriptome. (See Table S2).

**Extended Data 3.**
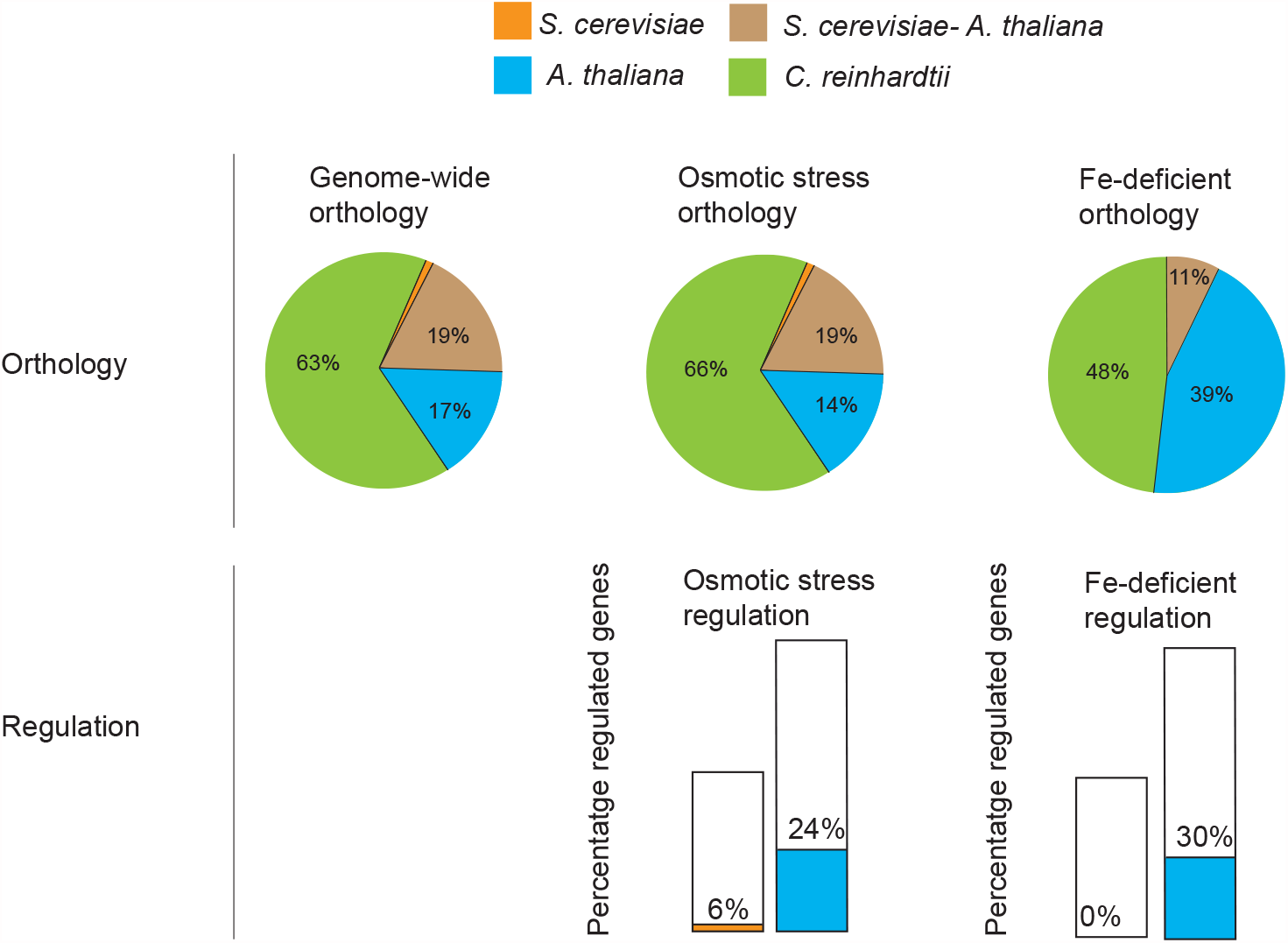
Conservation of the Chlamydomonas osmotic response across kingdoms. Pie charts representing the orthology of the Chlamydomonas genome with *S. cerevisiae* and *A. thaliana* genomes, orthology of differentially expressed genes in the Chlamydomonas osmotic time course and orthology of differentially expressed genes of Chlamydomonas iron-deficiency response; all othologies are based on reciprocal BLAST (> 1E-10) (See Material and Methods). Bars represent the percentage of genes differentially expressed upon osmotic stress or iron deficiency in Chlamydomonas with orthologous genes in *S. cerevisiae* (orange bars) or *A. thaliana* (blue) responding to the same stress^5,41,42^ (See Table S2).

**Extended Data 4.**
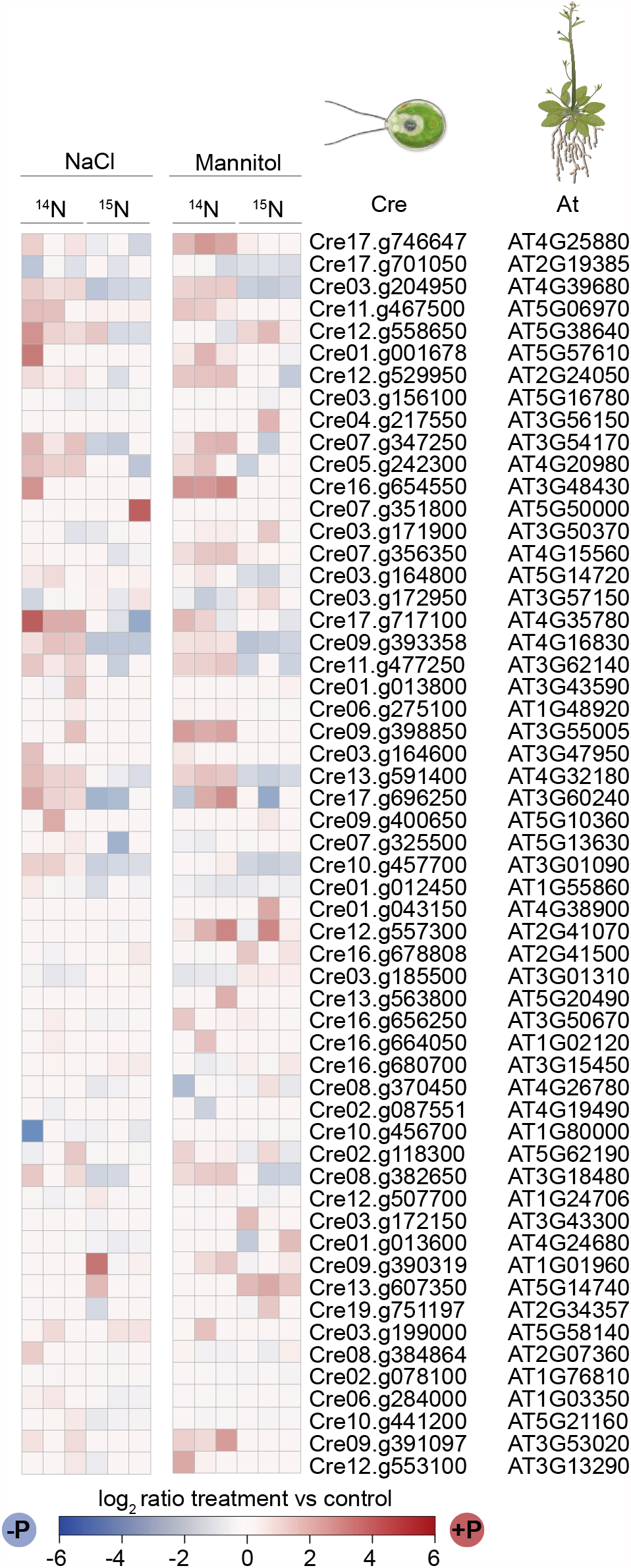
Phosphorylated proteins present in Chlamydomonas phosphoproteomic dataset and osmotically phosphorylated orthologues in Arabidopsis. Chlamydomonas peptides differentially phosphorylated (FC >1.5, FDR <0.01) with *A. thaliana* orthologous that have previously shown to be differentially phosphorylated upon osmotic conditions^43,44^ (See Table S3).

**Extended Data 5.**
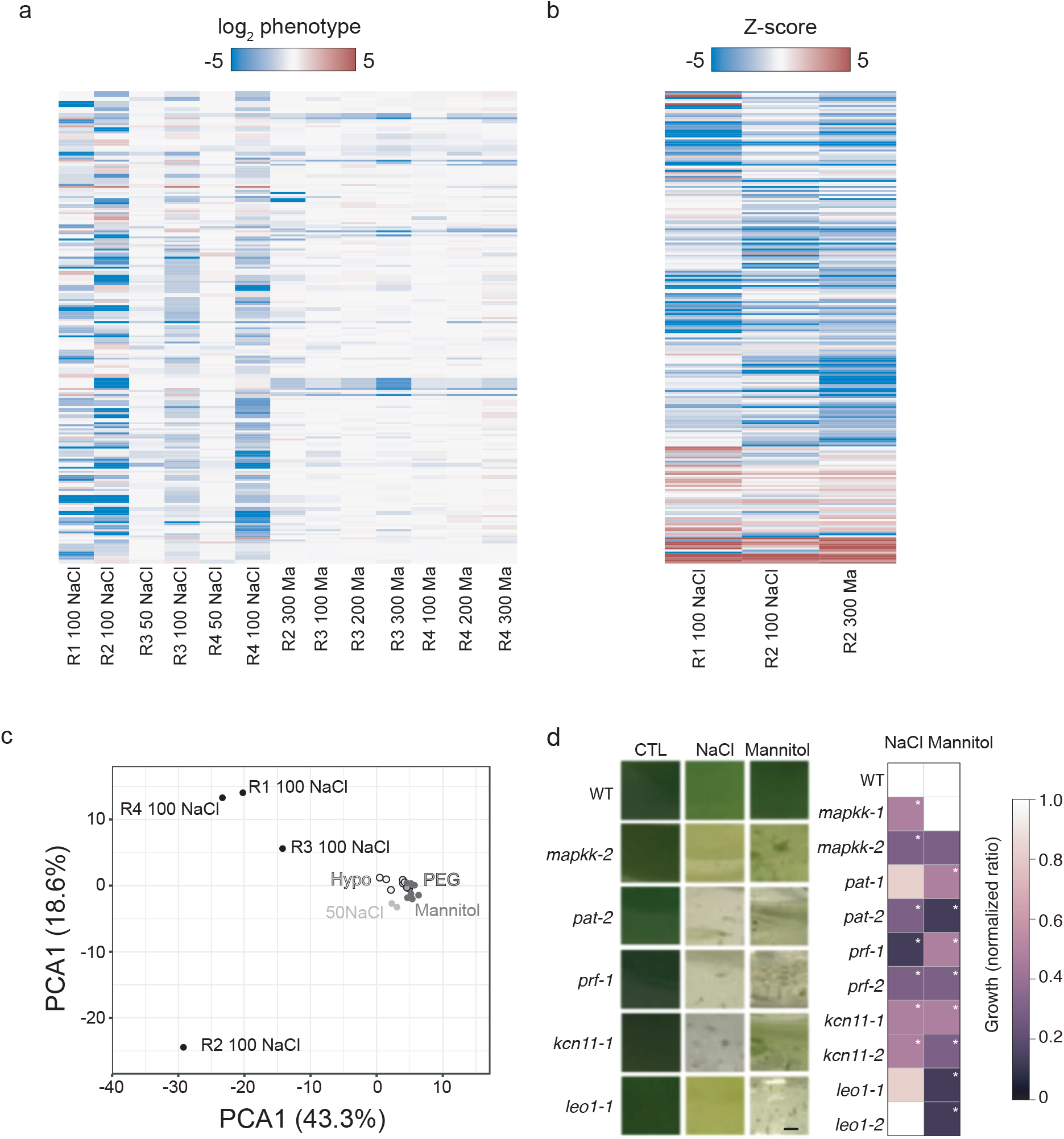
Validation of Chlamydomonas mutants with osmotic phenotypes. **a**, Median log2 phenotypes resulting from pooled mutant screens based on barcoded reads of hits selected for secondary screens (See Table S7). **b**, Z-score phenotypes of validated mutants. Z-score results from the quantification of chlorophyll RGB values (See Table S7). Note rows of mutants from A and B match horizontally. **c**, PCA of log2 median phenotype from different osmotic screens. **d**, Representative images of growth vessels 4 days post inoculation, grown in control media (CTL), 100 mM NaCl (NaCl) and 300 mM mannitol (Mannitol). Scale bar = 1 cm. (*Right*) Phenotype quantification 4 days post-inoculation under 100 mM NaCl (NaCl) and 300 mM mannitol (Mannitol) of selected Chlamydomonas hits. Growth of mutant strains upon stress were normalized to growth of wild type upon stress. * Represents p < 0.05 (two-way ANOVA).

**Extended Data 6.**
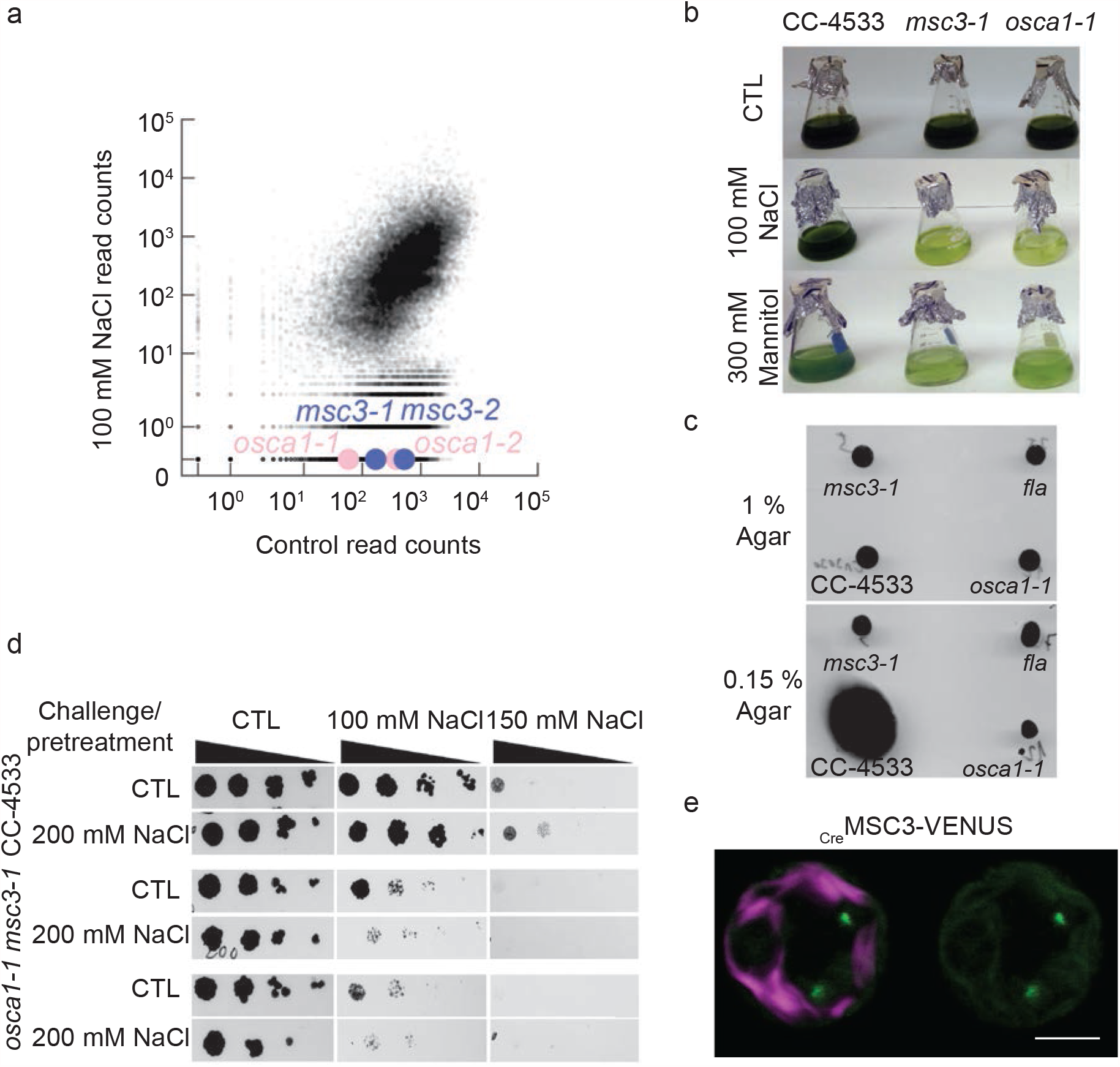
Functional conservation of osmoregulatory pathways across the green lineage. **a**, Scatter plot comparing barcode reads resulting from pooled screens in control condition and 100 mM NaCl treatment with *osca* and *msl* alleles highlighted. **b**, Growth of wild type (CC-4533) and *msc3-1* and *osca1-1* under control conditions (CTL), 100 mM NaCl and 300 mM mannitol four days after inoculation. **c**, Flagella phenotypes of *msc3-1* and *osca1-1* mutants. (*Top*) strains plated in 1% agar do not show colony differences, (*bottom*) strains plated in 0.15% agar show swimming differences based on colony size. As a positive control we used a flagella less mutant, *fla*. **d**, *msc3-1* and *osca1-1* mutants show acclimation defects to osmotic stress. Chlamydomonas CTL (CC-4533), *msc3-1* and *osca1-1* cells at the early mid-exponential phase (2E+06 cells/ml) were plated at 10-fold sequential dilutions (control raws, CTL) and Chlamydomonas cells at the exponential phase (2E+06 cells/ml) treated with 200 mM NaCL for 2 hours and plated in 10-fold dilutions (200 mM NaCl). **e**, Confocal image of a Chlamydomonas cell expressing MSC3 protein fused to VENUS fluorescent protein. Magenta signal corresponds to chloroplast autofluorescence and green fluorescent signal to VENUS.

**Extended Data 7.**
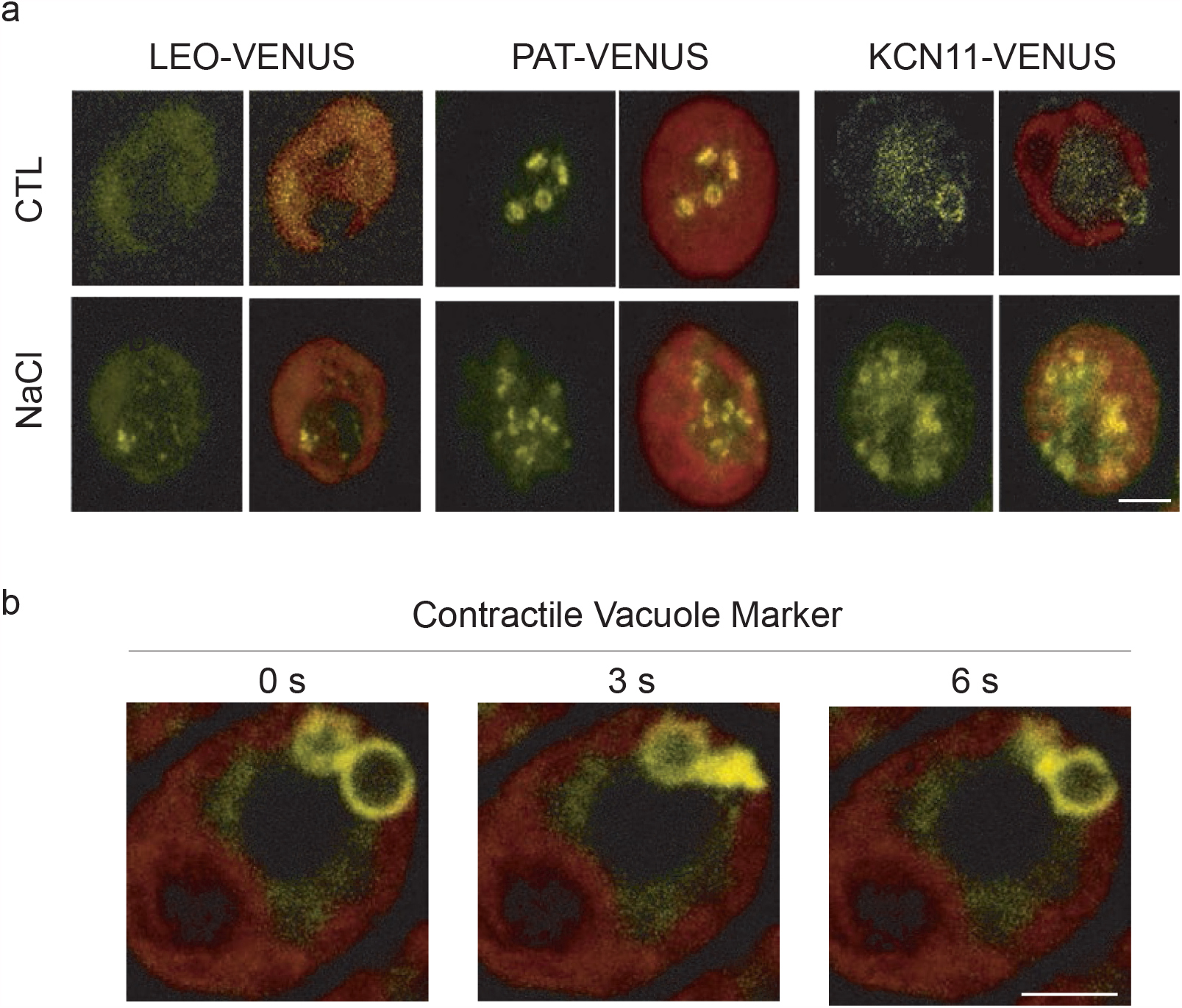
Protein localization upon osmotic stress of Chlamydomonas selected hits identified in the osmotic screens. **a**, Confocal images of Chlamydomonas cells expressing proteins fused to VENUS fluorescence protein under control conditions (CTL) and upon treatment with 100 mM NaCl (NaCl). Scale bar = 5 μm. **b**, Confocal images of Chlamydomonas cells expressing a contractile vacuole marker. Red color shows chloroplast autofluorescence and yellow VENUS signal. Scale bar = 5 μm.

**Extended Data 8.**
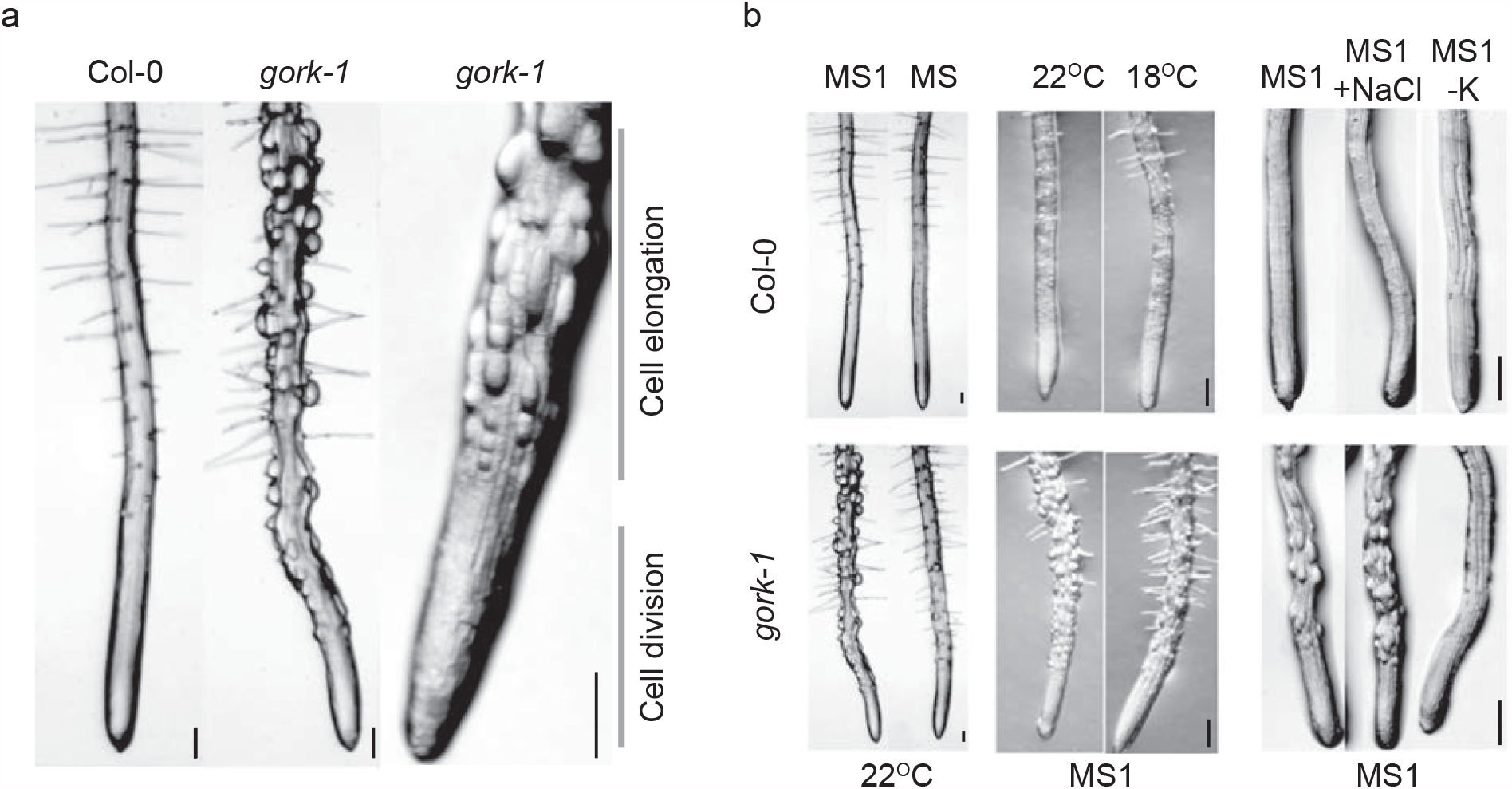
Growth dependent accumulation of potassium generates root hair inflated cells. **a**, Primary roots of 6 days-post germination wild-type (Col-0) and *gork-1* mutants. Scale bar = 100 μm. **b**, Primary root of 6 days-post germination Col-0 and *gork-1* mutants under different temperatures and sugar concentrations. MS1 media contains 1% sucrose, while MS does not contain sucrose. Scale bar = 100 μm. **c**, Primary root of 6 days-post germination Col-0 and *gork-1* mutants NaCl stress and media lacking potassium. MS1 media contains 1% sucrose, MS1 media supplemented with 140 mM NaCl (MS1, +NaCl), and MS1 media without any potassium (MS1 -K). Scale bar = 100 μm.

**Extended Data 9.**
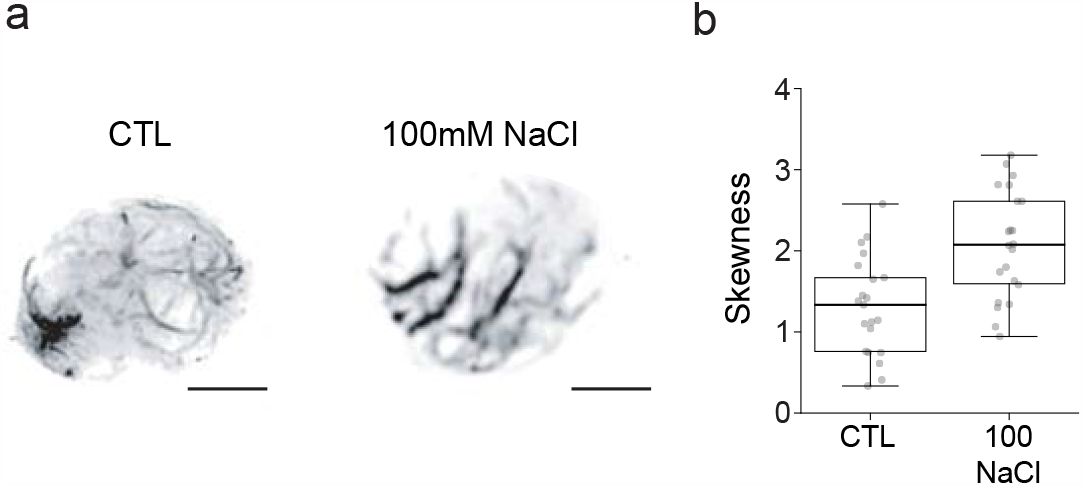
Salt stress promotes actin bundling in Chlamydomonas. **a**, Actin localization in Chlamydomonas cells under control conditions (CTL) and upon 16 hours treatment with 100 mM NaCl (NaCl). Actin filaments were visualized using Lifeact-NEOGREEN. Scale bar = 5 μm. **b**, Quantification of actin skewness in Chlamydomonas cells.

**Extended Data 10.**
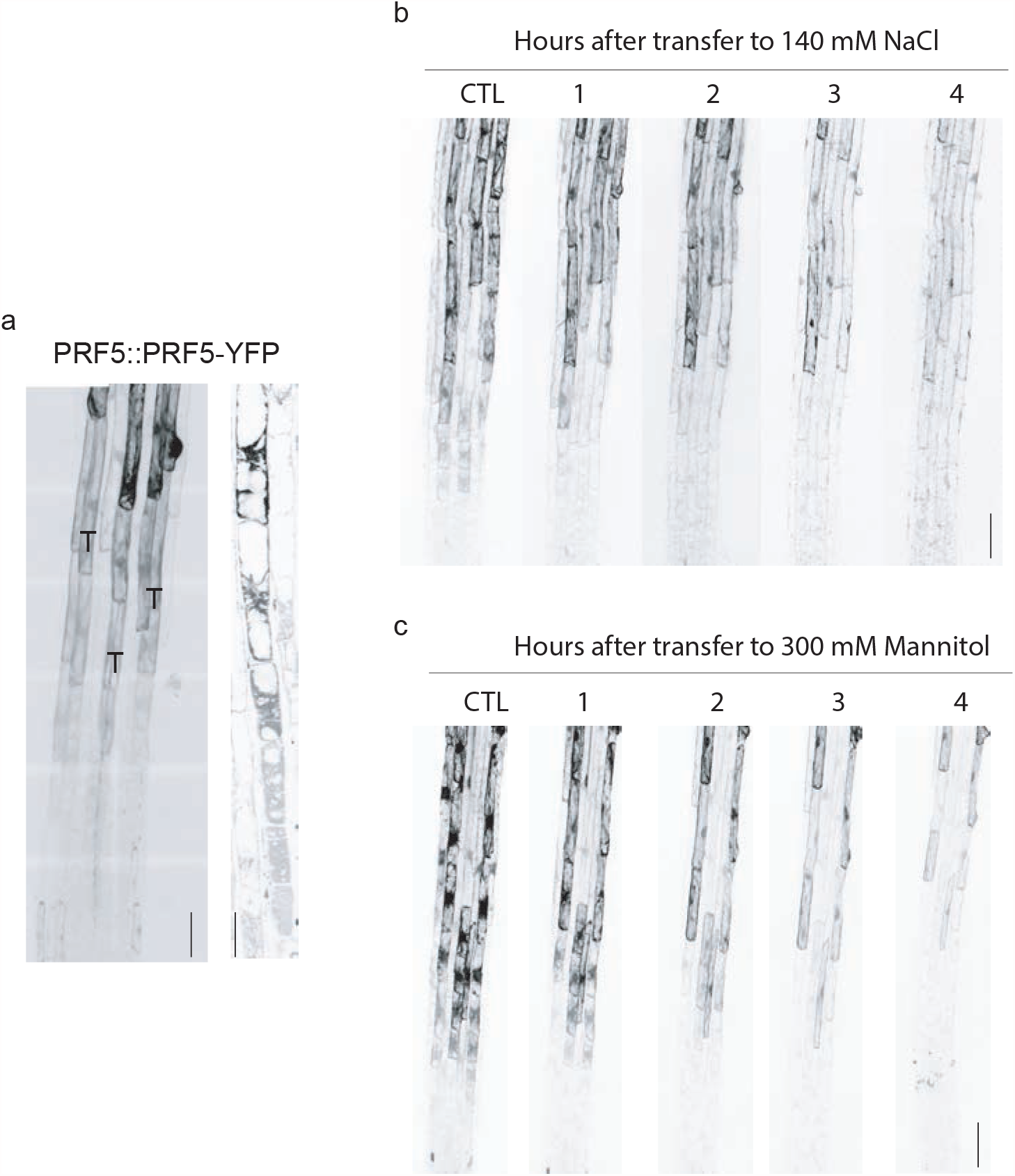
Profilin expression is restricted to root hairs and regulated by osmotic stress. **a**, Confocal image of Arabidopsis primary root elongation zone expressing ProPRF5::PRF5-YFP. T represents trichoblasts. Scale bar = 10 μm. **b**, Confocal images of Arabidopsis primary root elongation zone expressing ProPRF5::PRF5-YFP treated with 140 mM NaCl for the indicated hours. Scale bar= 20 μm. **c**, Confocal images of Arabidopsis primary root elongation zone expressing ProPRF5::PRF5-YFP treated with 300 mM mannitol for the indicated hours. Scale bar = 20 μm.

## List of supplementary materials

Table S1| Chlamydomonas RNAseq raw data

Table S2| Chlamydomonas transcriptomic analysis

Table S3| Chlamydomonas phosphoproteomics raw data

Table S4| Chlamydomonas phosphorylated hits, comparison with Arabidopsis

Table S5| Raw data osmotic mutant screens

Table S6| High confidence Chlamydomonas osmotic hits

Table S7| Chlamydomonas secondary screens

## Material and Methods

### General maintenance of *Chlamydomonas reinhardtii* strains and sample preparation

The background *Chlamydomonas reinhardtii* strain for all experiments was wild type (CC-4533). All *C. reinhardtii* strains were maintained in Tris-Acetate-Phosphate (TAP) solid media, 1.5% agar, with modified trace elements^45^ at 22 °C in low continuous light (∼15-10 μmol photons m^-2^ s^-^ _1_). Lines harboring VENUS-3xFlagged tagged genes were maintained in the same conditions with media supplemented with 20 µg/ml hygromycin. Routine maintenance intervals were every four weeks.

### RNAseq samples and analysis

RNA-seq samples: prior to sample collection wild-type strains (CC-4533) were refreshed in solid media and inoculated in liquid media, Tris-Acetate-Phosphate (TAP) with modified trace elements _45_ at 22 °C in low continuous light (100 μmol photons m-2 s-1). After 2 passages in liquid media (∼6 doublings/passage), cells growing in early-mid exponential phase (∼2E+06 cells/ml) were used for inoculation. Once cultures reached early-mid exponential phase (∼2E+06 cells/ml) different osmolyte concentrations were inoculated, 100 mM NaCl (Sigma) and 300 mM mannitol using 1 liter Erlenmeyer flasks and grown with mild agitation (∼150 rpm), 22 °C and low continuous light. Samples were collected at the designed timepoints from the same flask. Sample collection of ∼1E07 cells/ml was performed using a funnel with a POLYVINYLIDENE FLUORIDE membrane (GCWP4700, millipore) coupled to a vacuum station and subsequently cells were snap-frozen in liquid nitrogen and saved at -80 °C until processing.

For each treatment we performed 3 biological replicates with 2 control conditions, one at the beginning of the time course and another 24 hours after inoculation. RNA extraction was performed using Direct-zol RNA miniprep kit (R2050)) following manufacturer’s instructions including in-column DNase I treatment prior RNA washing and elution steps. Each RNA sample was run on an Agilent 2100 Bioanalyzer RNA 6000 Nano chip for quantification and quality control. RNA samples were submitted to the Stanford Functional Genomics facility for subsequent library preparation and sequencing. RNA-seq libraries were made using the Kapa mRNA HyperPrep kit (Roche, KK8540) following manufacturer’s protocols. Libraries were pooled based on fragment analyzer concentrations. Sequencing was performed on Nextseq high-output flow cell, 1×75 bp run (Ilumina).

RNA-seq reads were aligned using BWA to the Chlamydomonas reference genome 5.6 ^46^. DESeq2 package was used to call transcripts as differentially expressed at a false-discovery-rate (Benjamini-Hochberg method) FC >2, (FDR <0.01). GO-term enrichment analysis was performed with the BINGO plugin for Cytoscape^47^. To perform GO-term-enrichment analysis of differentially regulated transcripts, we used several approaches. First, we used GO terms from the Joint Genome Institute (JGI) for Chlamydomonas genome 5.6 version, and used these as a reference set. Second, we used the annotations of the homologous genes in *Arabidopsis thaliana*. To identify these homologues, we identified gene pairs by reciprocal best BLAST hit (RBH) and selected “Best hit” for each pair having the lowest e-value (Onishi 2018), setting an e-value threshold of 1E-10.

### Proline quantification

Proline quantification in Chlamydomonas cells was adapted from^48^. Briefly, 1E05 cells from control or treated samples in early-mid exponential growing phase were collected and transferred to 96 wells plates together with L-proline standards. Subsequently the same volume of glacial acetic acid and acid ninhydrin was added to each well, sealed and incubated for 1 hour at 100 °C. Following incubation and cool down of samples chromophore was extracted with Toluene and the absorbance read at 520 nm using toluene as blank. Determination of proline concentration was performed from a standard curve.

### Phosphoproteomics samples and analysis

Phosphoproteomics samples: prior to sample collection wild-type strains (CMJ030/CC-4533) were refreshed in solid media and inoculated in liquid media, Tris-Acetate-Phosphate (TAP) with modified trace elements ^45^ at 22 °C in low continuous light (100 μmol photons m-2 s-1). After 2 passages of cells growing in early-mid exponential phase (∼6 doublings/passage), with either 14NH_4_Cl or 15NH_4_Cl as a sole source of nitrogen, cells growing in early-mid exponential phase (∼2E+06 cells/ml) were used to inoculate 2-liter bottles with bubbled air and constant stirring at an initial concentration of 2E04 cells/ml. This ensured 15N growth for at least eight generations. Cells at early-mid exponential phase were spun out (2,000g, 5 min, 4 °C) and resuspended in a 1:1 (v/w) ratio of ice cold homogenization buffer (290 mM sucrose, 250 mM Tris-HCL (pH = 8), 25 mM EDTA, 25 mM sodium fluoride, 50 mM sodium pyrophosphate, 1 mM ammonium molybdate, 0.5% polyvinyl pyrovinyl pyrrolidone in water and 1 cOmplete EDTA-free protease inhibitor (Sigma-Aldrich)/50 ml). The cell slurry was then added drop wise to liquid nitrogen to form small Chlamydomonas pellets approximately 5 mm in diameter. Samples were stored at -80 °C.

Samples for phosphopeptide enrichment were prepared as previously described (Minkoff 2014) with minor modifications. Samples were combined in an experimental pair consisting of one treated sample grown in 14NH_4_Cl and one control sample growing in 15NH_4_Cl. For the second reciprocal experimental pair, the samples were combined in the inverse fashion (15NH_4_Cl treated and 14NH_4_Cl control). Frozen cells were combined at 1:1 weight ratio prior homogenization, with a total weight of 4 grams (2 grams of cells grown in 14NH_4_Cl and 2 grams of 15NH_4_CL). Three biological replicates were processed for each treatment condition. For all experiments, samples were processed in homogenization buffer supplemented with phosphatase inhibitors using a sonicator (1 cm probe, (12 × 5 seconds) x2, and 50% duty cycle (Bransom 45 Digital Sonifier (Marshall Scientific)) while kept on ice. The resulting homogenate was filtered through two layers of Miracloth (Calbiochem) and spun out 15 minutes at 1,500 g and 4 °C. Subsequent steps were performed as described previously^49^.

Phosphopeptide-enriched samples were analyzed on an LTQ-Orbitrap XL mass spectrometer (Thermo Scientific). Acquired data files containing MS/MS spectra were searched against the JGI *Chlamydomonas reinhardt* annotation v5.6 protein database using MASCOT software (Matrix Science). Searches were performed using settings for both 14N and 15 N protein masses. MASCOT search results were filtered to maintain a 1% false discovery rate at the peptide level using a reverse-protein sequence database. Quantitative ratio measurements from peak areas were performed using Census software. Only phosphopeptides showing reciprocal changes of 2-fold or greater in two of the three replicates were selected for data shown in Figure 1. Phosphopeptides showing reciprocal changes of 1.5-fold or greater were used to identify homologous phosphorylated proteins in Arabidopsis (Extended Data 3).

### Chlamydomonas mutant screen

Library maintenance, pooling, competitive growth, DNA extraction, barcode amplification and library preparation were performed as described^50^. We used several approaches to identify mutants with growth defects, as described^50^. High-confidence gene-phenotype relationships (Figure 2, Extended Data 6, Table S4) were based on false discovery rate (FDR <0.3) performed on p-values of genes with more than 2 alleles using the Benjamini-Hochberg method^51^.

### Secondary screens

Chlamydomonas mutants selected for secondary screens were picked from the Chlamydomonas mutant library and inoculated in 96-well plates containing liquid TAP and 10 µg/ml paromomycin. Plates were re-inoculated 3 times (ensuring ∼10 doublings), and early-mid exponential cells were used to inoculate control plates (TAP liquid), plates supplemented with 100 mM NaCl or plates containing 300 mM mannitol. Plates were grown under low light conditions. 5 days post-inoculation plates were scanned using a CanonScan 9000F flatbed scanner (Canon). Images were quantified using Fiji. 8-bit RGB images were converted to gray scale and average pixel intensity was measured with a sampling area including the entire well. Wells with bubbles and/or irregular clamping were avoided and not quantified. Comparisons between average pixel intensity were used to estimate the mutant growth compared to wild type. Each plate contained 2 biological replicates of each mutant and multiple wild-type colonies to avoid any positional effect. The resulting Z-scores can be found in Table S7.

### Chlamydomonas protein localization

To generate fluorescently tagged proteins in Chlamydomonas, we used pRAM118 (a generous gift from Silvia Ramundo). Open reading frames were PCR amplified (Phusion Hotstart II polymerase, ThermoFisher Scientific or KOD DNA polymerase, Millipore) from genomic DNA or BAC, gel purified (MinElute Gel Extraction Kit, QIAGEN) and cloned in frame with the C-terminal VENUS-3xFLAG tag by Gibson assembly. Primers were designed to amplify target genes from their predicted start codon up to, but not including the stop codon. All resulting constructs were verified by Sanger sequencing. Constructs were linearized by EcoRV prior to transformation. Electroporation was performed with cells grown to ∼8E06 cells/ml, resuspended at ∼8E08 cells/ml in CHES buffer at room temperature, and electroporated in a volume of 125 µl in a 2-mm-gap electro cuvette using a NEPA21 square-pulse electroporator (Bulldog Bio), using two poring pulses of 250 and 150 V for 8 msec each, and five transfer pulses of 50 msec each starting at 20 V with a “decay rate” of 40% ^52^. Cells were transferred to a 15-ml centrifugation tube containing 8 ml TAP plus 40 mM sucrose. After overnight incubation at 24 °C under low light, cells were collected by centrifugation and spread on TAP agar (1.5% w/v) plates containing 25 µg/ml hygromycin.

Colonies for plate reader analysis resulting from transformation plates were picked and transferred to a 96-well microplate plate containing 150 µl liquid of TAP supplemented with 25 µg/ml hygromycin, without shaking. Fluorescence readings were acquired with excitation and emission wavelengths of 515 and 550 nm, TECAN infinite 200 Pro microplate reader. Five independent colonies with values greater than background were further selected for imaging. Expression of Cre12.g524950-VENUS (LEO) was further confirmed with Western blot.

### Arabidopsis growth and treatments

Arabidopsis thaliana Col-0 ecotype was used in this study. A complete list of the mutant lines used in this study can be found Table S9. Seeds were surface sterilized by washing with 20% bleach for 5 minutes and rinsed with sterile deionized water four times. After stratification seeds were grown in 10 × 10 cm petri dish plates containing sterile full MS media (MSP01-50LT; Caisson), 1% Sucrose (Sigma-Aldrich), 0.7% Gelzan, 0.05% MES (Sigma-Aldrich) adjusted to pH = 5.7 using 1 M KOH. Screening of Arabidopsis mutants was performed with seedlings growing under standard media and transferred to plates containing 140 mM NaCl (Sigma-Aldrich) or 300 mM mannitol (Sigma-Aldrich) after 4 days post germination.

Growth of seedlings was performed in a Percival CU41L4 incubator at constant temperature 24 °C with 14 h light and 10 h dark cycles at 130 μmol m^2^ s^-1^ light intensity. Plates were sealed using micropore tape (3 M). Plates were placed vertically to allow vertical growth of the roots.

Images of seedlings were captured using Epson perfection V800 Photo color scanner, root length was quantified using Fiji^53^.

### Microcopy

Confocal laser microscopy was performed on a Leica TCS SP8 inverted confocal scanning microscope in resonant scanning mode with LASX software. Chlamydomonas cells were mounted in chambered coverglass (Ibidi, 80826), covered with 2% low melting agar and imaged with 93x/1.3 NA glycerin immersion objective. Arabidopsis roots were mounted in chambered coverglass (ThermoFisher, 155360) with a pad of Gelzan MS media or gelzan MS media supplemented with corresponding treatment. Root growth time lapse movies were taken using a 20x/0.75 NA glycerin immersion objective. For each root 5 tiles were taken using the Navigator mode, with the root tip position in the middle of the second tile at the beginning of the time course, to ensure capture of the root growth upon the 16 hour period. Fluorescence of LTi6b-YFP, ABD2-GFP and GCaMP6 lines was captured with a white light laser, excitation set at 488 nm and detection from 500-550 nm with a HyD SMD hybrid detector (Leica).

All image quantifications were performed with Fiji^53^. Quantification of root cell death was performed using Z-stack projections and manual counting of cell death. Actin quantification was performed with stack images taken from the first visual cortical actin in the epidermal elongating cells, collecting 20 steps of 1 μm each. Imaging parameters gain and pinhole were selected such that individual actin filaments could be observed, but actin filament bundles were not saturated. All cropped images used for quantification were taken from original 8-bit files. Actin images were cropped along the entire length of every cell in the elongation zone. Skewness was analyzed according to^54^. No image processing was applied to maximum-intensity projections that were analyzed for skewness. The size of each cell crop was maintained constant across all images and was smaller than the entire length of a cell (10 × 10 μm). Relative angle of actin filaments was computed using Fiji and aligning cells horizontally. Calcium spikes quantification was performed at the transition zone in root epidermal cells (∼200 µm from the stem cell niche) during a period of 4 hours. Registration was applied to time lapses using the HyperStackReg plugin. A MaxEntropy Threshold was applied, and 8 ROIs (24×24 µm) were selected across a 150 µm area to measure the signal corresponding to calcium spikes. A minimum of 6 pixels was used to exclude sporadic signals. Calcium spikes and their duration were confirmed manually.

## Data availability

Strains, plasmids and plasmid sequences are available upon request and will be deposited at the Chlamydomonas Resource Centre.

